# Increased spatial coupling of integrin and collagen IV in the immunoresistant clear cell renal cell carcinoma tumor microenvironment

**DOI:** 10.1101/2023.11.16.567457

**Authors:** Alex C Soupir, Mitchell T Hayes, Taylor C Peak, Oscar Ospina, Nicholas H Chakiryan, Anders E Berglund, Paul A Stewart, Jonathan Nguyen, Carlos Moran Segura, Natasha L. Francis, Paola M. Ramos Echevarria, Jad Chahoud, Roger Li, Kenneth Y. Tsai, Jodi A. Balasi, Yamila Caraballo Peres, Jasreman Dhillon, Lindsey A. Martinez, Warren E. Gloria, Nathan Schurman, Sean Kim, Mark Gregory, James Mulé, Brooke L Fridley, Brandon J Manley

## Abstract

**Background:** Immunotherapy (IO) has improved survival for patients with advanced clear cell renal cell carcinoma (ccRCC), but resistance to therapy develops in most patients. We use cellular-resolution spatial transcriptomics in patients with IO naïve and IO exposed primary ccRCC tumors to better understand IO resistance. Spatial molecular imaging (SMI) was obtained for tumor and adjacent stroma samples. Spatial gene set enrichment analysis (GSEA) and autocorrelation (coupling with high expression) of ligand-receptor transcript pairs were assessed. Multiplex immunofluorescence (mIF) validation was used for significant autocorrelative findings and the cancer genome atlas (TCGA) and the clinical proteomic tumor analysis consortium (CPTAC) databases were queried to assess bulk RNA expression and proteomic correlates.

**Results:** 21 patient samples underwent SMI. Viable tumors following IO harbored more stromal CD8+ T cells and neutrophils than IO naïve tumors. *YES1* was significantly upregulated in IO exposed tumor cells. The epithelial-mesenchymal transition pathway was enriched on spatial GSEA and the associated transcript pair *COL4A1*-*ITGAV* had significantly higher autocorrelation in the stroma. Fibroblasts, tumor cells, and endothelium had the relative highest expression. More integrin αV+ cells were seen in IO exposed stroma on mIF validation. Compared to other cancers in TCGA, ccRCC tumors have the highest expression of both *COL4A1* and *ITGAV*. In CPTAC, collagen IV protein was more abundant in advanced stages of disease.

**Conclusions:** On spatial transcriptomics, *COL4A1* and *ITGAV* were more autocorrelated in IO-exposed stroma compared to IO-naïve tumors, with high expression amongst fibroblasts, tumor cells, and endothelium. Integrin represents a potential therapeutic target in IO treated ccRCC.

## Background

The use of immunotherapy (IO) in the treatment of clear cell renal cell carcinoma (ccRCC) has improved response rates and survival outcomes compared to those of the targeted therapy era. ^1–3^ However, IO resistance limits progression free survival to 11-15 months. ^1, 2^ Certain histological features of ccRCC, such as sarcomatoid morphology, portend an excellent response to IO treatment. ^4^ Yet reliable biomarkers of IO responsiveness or resistance in other cancers have not been shown to translate to ccRCC.^5^ Thus, there is an enormous clinical need to discover distinct mechanisms that stratify IO responsive from IO resistant phenotypes.

Elements of the tumor immune microenvironment (TIME) have been found to fluctuate as patients with ccRCC progress. ^6, 7^ Large scale efforts to characterize the ccRCC TIME through bulk transcriptomic sequencing provided an understanding of the macro-level immune cell composition within ccRCC. ^8, 9^ These studies have revealed the association of M2-macrophage infiltration, T cell exhaustion, and angiogenesis enriched molecular profiles with survival outcomes and treatment responses in both the tyrosine kinase inhibitor (TKI) and IO eras. Advances in single cell RNA sequencing (scRNA-seq) allowed profiling of heterogenous cell populations within tumor and the surrounding ‘normal’ tissue. ^10^ More recently this technology has been employed to identify differences in various T cell populations between IO responsive and IO resistant tumors.^11^ In these ways, scRNA-seq has deepened our understanding of various cell lineages amongst various ccRCC tumor stages; however, many questions remain regarding the biology of cell-cell interactions across the clinical spectrum in ccRCC. A limitation of scRNA-seq is that the tissue is disassociated and thus does not preserve the tissue architecture (i.e., removes the spatial locations and relationships between individual cells).

Technological advancements may overcome some of the disadvantages of bulk and single cell methods. Specifically, spatial proteomics has identified the association of certain myeloid cell line clustering within tumoral regions and poor treatment response with immunotherapy (IO), as well as survival outcomes. ^6, 11, 12^ These studies utilized spot-based or mini-bulk resolution but does not elucidate cell-cell crosstalk. Recently, high-plex spatial transcriptomics has allowed simultaneous analysis of cell-level variations in gene expression on a single slide. In situ hybridization of RNA transcripts in formalin fixed specimens can yield subcellular spatial information and recapitulates the signals seen in bulk RNA sequencing and scRNA-seq.^13^ On some platforms, this allows investigation of nearly 1000 transcripts amongst millions of cells on the same slide.^14^

Utilizing the NanoString CosMx Spatial Molecular Imager (SMI) for spatial transcriptomic analysis,^14^ we sought to evaluate the TIME in three clinically relevant patient populations and TIME of their ccRCC tumors: those who had yet to undergo IO treatment and had tumors without sarcomatoid features (i.e., IO naïve non-sarcomatoid ccRCC tumors), those who had yet to undergo IO treatment and had tumors with sarcomatoid features (i.e., IO naïve sarcomatoid ccRCC tumors), and those treated with IO (i.e., IO exposed ccRCC tumors). These three patient cohorts represent clinically diverse outcomes, with patient’s tumors that are IO naïve and patient’s tumors that have sarcomatoid features having potential enhanced initial response to IO treatment. Residual tumors present after IO treatment have sustained eco-evolutionary selective pressures and subsequently contain subpopulations of IO resistant tumor cells that harbor the genomic potential for clinical progression. ^15, 16^ Thus, our study objective was to define the unique cellular and spatial characteristics of these clinically relevant ccRCC cohorts utilizing high resolution spatial transcriptomic technology.

## Results

### Quality Control

Each patient tumor sample had a 1mm sample obtained from a region of tumor and stroma near the respective tumor-stroma interface. SMI was obtained on three tissue microarrays (TMAs) using the human Immuno-oncology panel (consisting of 978 RNA probes). Between stromal and tumor primary disease fields of view (FOVs), 40 FOVs passed the applied quality filters (See Methods, **Table 1**, **Supplementary Table 1**). The number of cells on remaining FOVs ranged from 930 to 6090, with a median of 2969 cells per FOV.

**Table 1:**
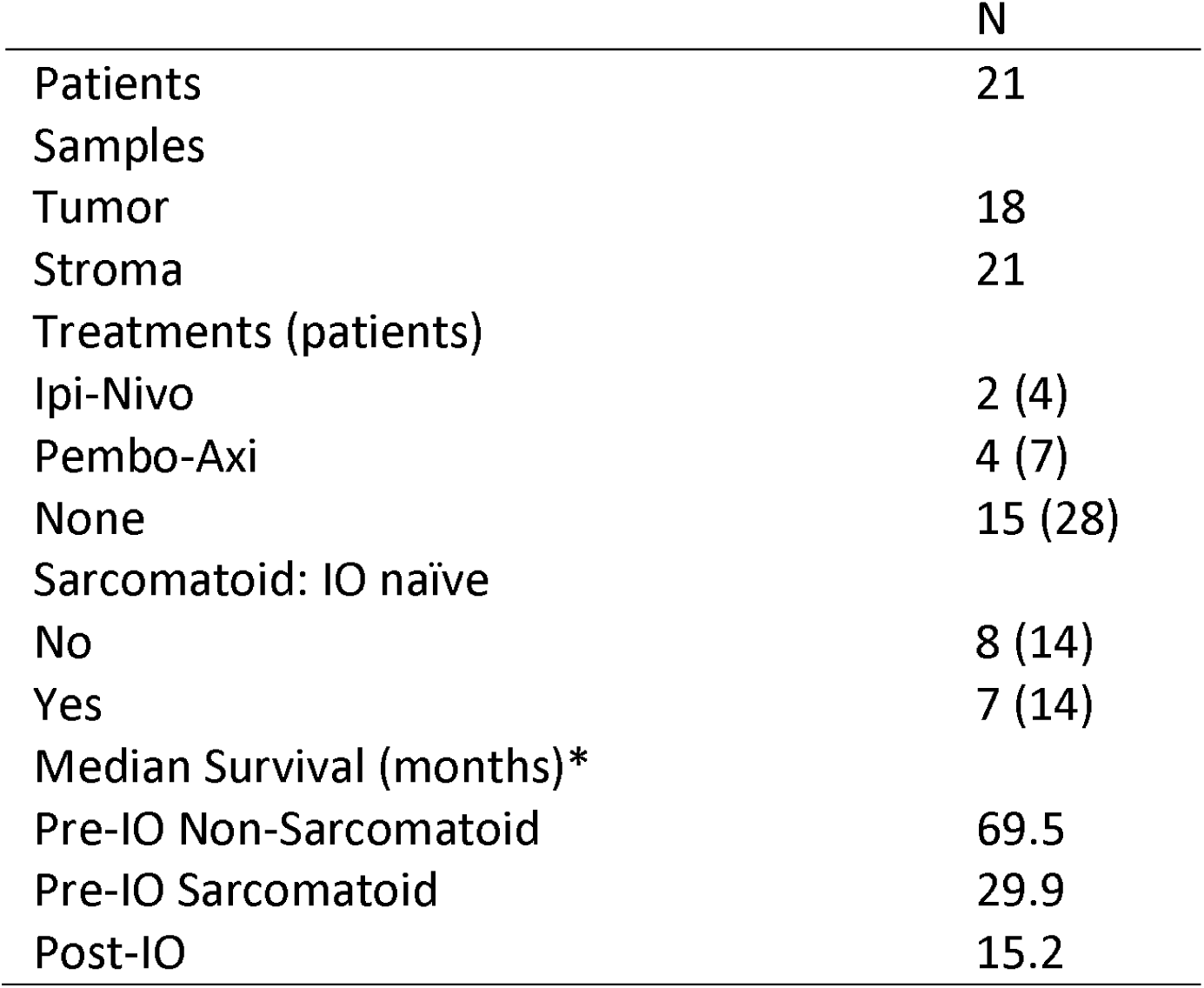
Patient cohorts for this study.

### Subclustering T cells and Mononucleic Phagocytes (MNPs)

Subclustering for T cells (**Figure 1A**) with Louvain clustering identified 8 unique clusters from the first 50 principal components (**Figure 1B**). Between these 8 unique subclusters, 112 unique genes were identified as differentially expressed in at least one cluster (absolute log fold change [LFC] > 0.25 and p < 0.004) (**Supplementary Table 2**). One of the new Louvain clusters was dominated by CD4+ T cells and higher *FOXP3* gene expression, and thus it was identified as regulatory T cells (**Figure 1B**). Subclustering of MNPs resulted in 14 distinct clusters (**Figure1C-D**). Among the 14 clusters, 189 unique genes were found to be differentially expressed (absolute LFC > 0.25 and p < 0.004; **Supplementary Table 2**). Seven of those clusters showed signatures of MNPs while the other six were more consistent with regulatory T cells, collecting duct cells (x3), neutrophils (x2), and B cells (**Supplementary Table 2; Figure 1D**). **Figure 1E** shows uniform manifold approximation and projection (UMAP) of all cells with their final phenotype assignments.

**Figure 1:**
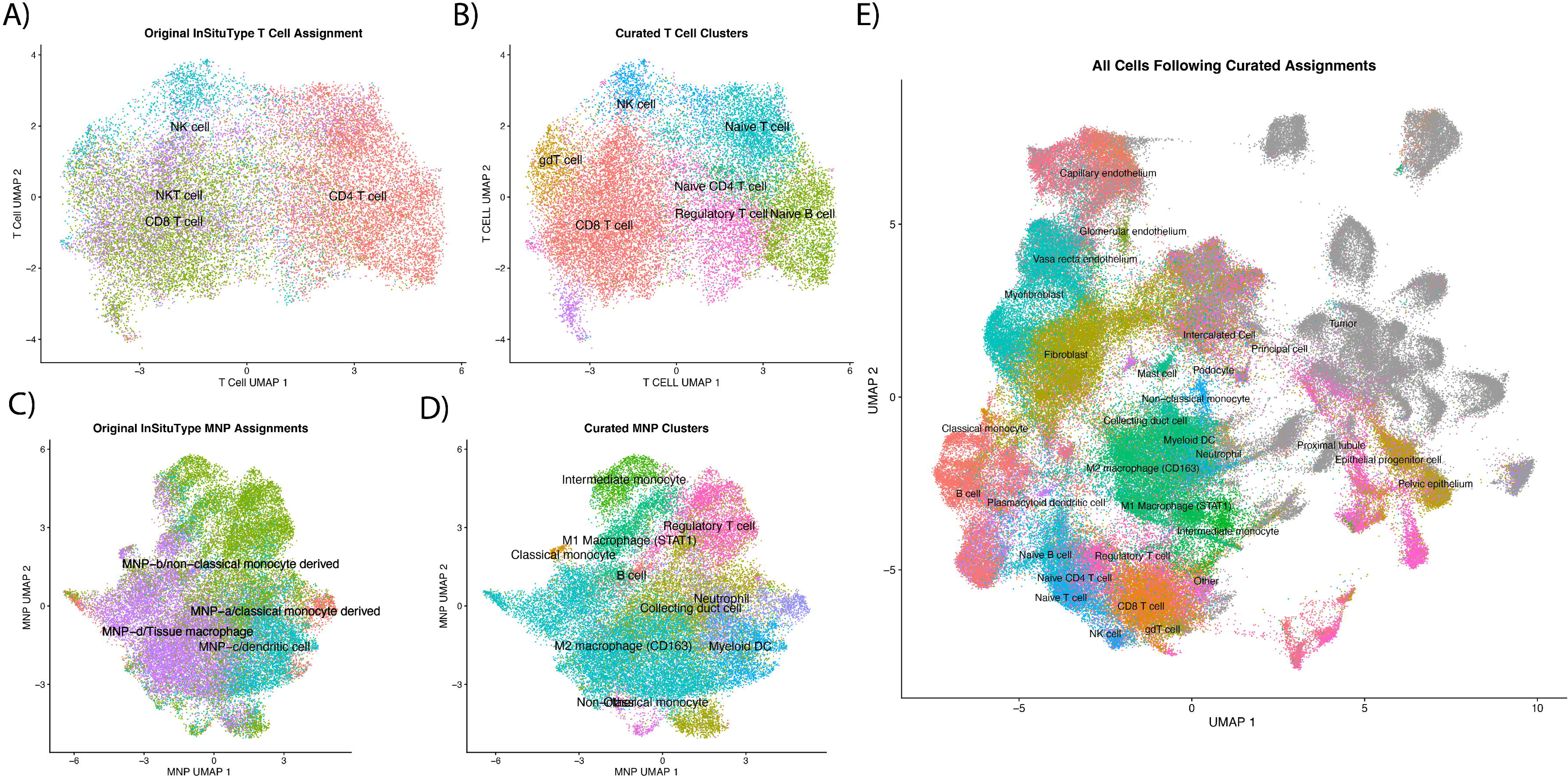
Phenotyping of cells following assignment with ‘InSituType’. A) and C) show UMAPs of T cells and mononucleic phagocytes (MNP), respectively, calculated with all cells in all fields of view (FOVs). Refined phenotypes from calculating new PCA/UMAP, clustering the subset with Louvain, and identifying markers with ‘FindAllMarkers’ can be seen in B) and D). Final cell assignments of all cells, including tumor cells, are shown in E).

### Identifying Malignant ccRCC Cells

UMAPs were created (**Figure 2A and 2F**) and gene expression of *TP53*, *EGFR*, *MYC*, and *VEGFA* plotted (**Figure 2B and 2G**) to identify FOVs for classifying tumor and normal cells, respectively. As anticipated, *VEGFA* was high for proximal tubule cells on tumors FOVs (**Figure 2C**) while stromal FOVs showed lower expression in proximal tubule cells (**Figure 2H**). The spatial context of these cells can be further seen in their H&E images (**Figure 2D, 2I**) as well as the cell typed polygon plots (**Figure 2E, 2J**).

**Figure 2:**
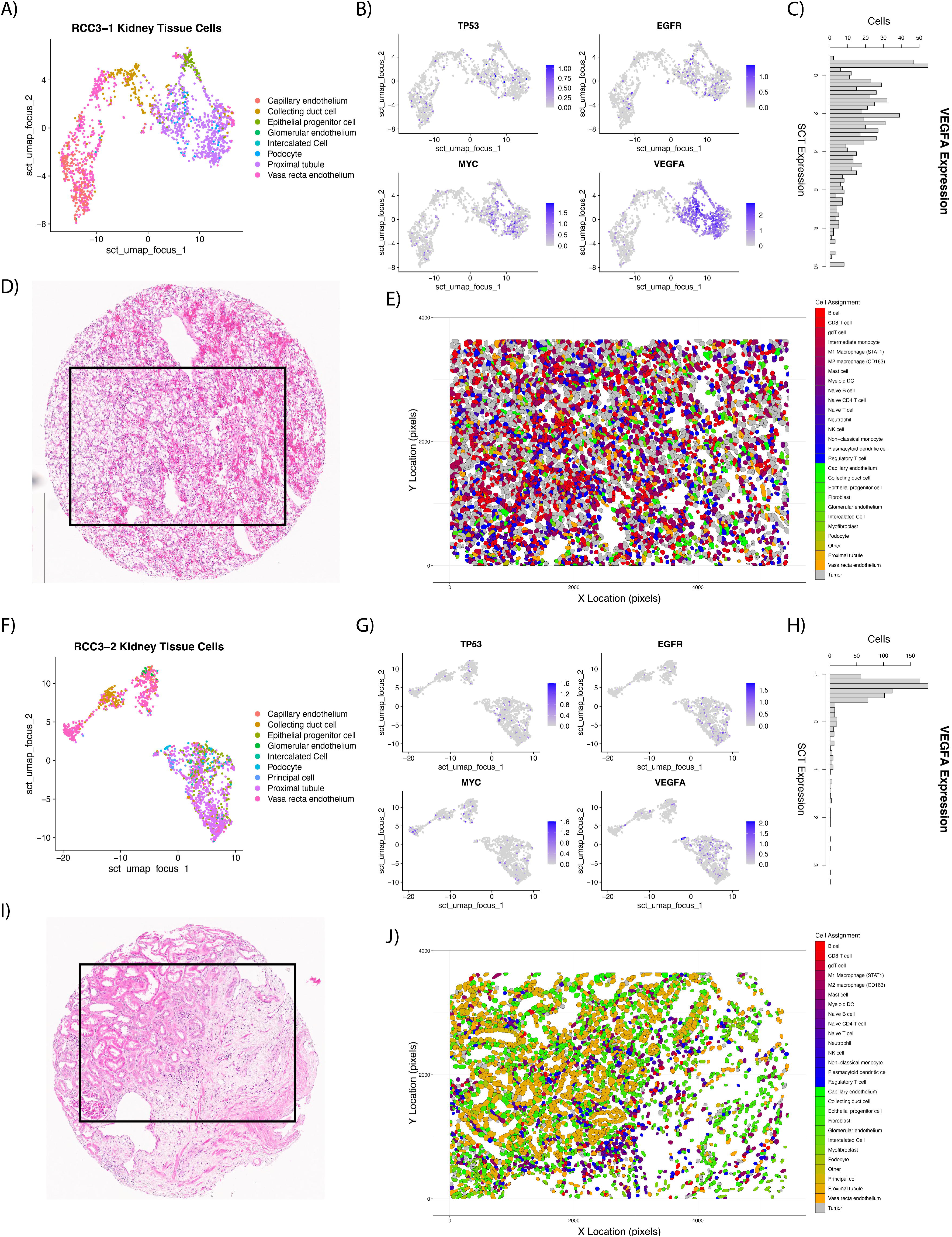
Examples of FOVs used for malignant cell identification with LASSO generalized linear models. UMAPs were created of kidney tissue specific cells (non-immune, non-fibroblast) for a tumor (A) and stroma (F) FOV. Gene expression was plotted over the UMAP for TP53, EGFR, MYC, and VEGFA with ‘FeaturePlot’ to aid in identifying malignant cells (B and G). VEGFA showed high expression in malignant proximal tubule cells (C) while expression in normal proximal tubule cells (H) was low. H&E images (D and I) who cores sent for CosMx SMI. After ‘InSituType’ and malignant cell classification, polygon plots were constructed with final cell assignments (E and J).

Differential gene expression was performed to find marker genes specific to either the malignant or normal proximal tubule cells, resulting in 51 differently expressed genes (**Supplemental Table 3**). Gene expression of these 51 genes were passed to a logistic regression model with Least Absolute Shrinkage and Selection Operator (LASSO, 10-fold cross validation) along with the class of the proximal tubule cell (normal or malignant) to shrink the number of genes needed to predict class. This decreased the number of genes from 51 to 43 (**Supplementary Table 3**) to be used within the generalized linear model to predict malignant classification (**Figure 2E, 2J**). Further review of the LASSO predictions by a genitourinary pathologist (JD), led to only one FOV reclassification for downstream analyses to ‘stroma’ (FOV17 on slide RCC4). Average cohort abundance of tumor/stroma is shown in **Supplementary Figure 1**.

### Higher abundance of CD8 T cells and neutrophils in the IO exposed TIME and M2 macrophages in the sarcomatoid TIME

Within the tumor FOVs, there were no difference in phenotype abundances between IO naïve and IO exposed tissue samples (**Supplementary Table 4**). However, the cell phenotypes in the stromal FOVs showed CD8+ T cells (false discover rate [FDR] = 0.0007) and neutrophils (FDR = 0.011) significantly higher in the IO exposed samples compared to IO naïve (**Supplementary Table 4**). Non-malignant proximal tubule cells were significantly more abundant in IO naïve FOVs (FDR = 0.018) while tumor cell counts were not significantly different between IO exposed and IO naïve FOVs (FDR = 0.939). Myofibroblasts and fibroblasts were not significantly more abundant in the stromal FOVs of IO naïve than IO exposed (FDR= 0.637 and 0.503, respectively).

Comparing IO naïve samples with and without sarcomatoid features, none of the cell phenotypes showed differences in abundances in tumor FOVs (**Supplementary Table 4**). In the stromal FOVs, M2 macrophages showed significantly higher abundance in sarcomatoid samples compared to non-sarcomatoid samples (FDR = 0.044, **Supplementary Table 4**).

### Minimal changes in cell clustering differences between patient cohorts

We used a quantitative framework leveraging Ripley’s K estimates, a measure of spatial heterogeneity commonly used in ecology and economics, to identify differences in cell type spatial clustering. ^17^ We observed that principal cells in the stroma of those exposed to IO were significantly more clustered than they were in IO naïve samples (FDR = 0.046; **Supplementary Table 5**). All other cell types showed no differences between IO naïve, and IO exposed.

The Ripley’s *K* estimates of individual cell types in IO naïve non-sarcomatoid and sarcomatoid feature tissues did not show significant differences in either tumor or stromal FOVs (**Supplementary Table 5**).

### *YES1* significantly upregulated in tumor cells following IO exposure

Pseudo-bulk analysis is the aggregated expression of genes over all cell types utilizing single cell data. ^18^ On pseudo-bulk analysis, IO exposed tumor FOVs showed 13 genes significantly downregulated and a single gene with higher expression than IO naïve (*YES1*, FDR 0.084; **Supplementary Table 6)**. The most significant downregulated gene in IO exposed tumor FOVs was *TPSB2* (FDR = 0.019).

In our differential expression analysis by cell phenotype, tumor cells from IO exposed tumor FOVs only showed 15 genes significantly higher than tumor cells from FOVs naïve to IO, *YES1* was the most significant (FDR = 0.026). In the IO exposed FOVs, principal cells and pelvic epithelium had the largest number of genes significantly higher (633 and 113 genes, respectively; **Supplementary Table 7**).

On pseudo-bulk analysis of IO naïve sarcomatoid vs non-sarcomatoid samples, there were no significant different expressed genes (**Supplementary Table 7**). In sarcomatoid tumor FOVs, differential expression by phenotype revealed tumor cells having significant downregulation of *IL17A* compared to non-sarcomatoid tumor FOVs (FDR < 0.001) along with seven other genes. No genes were significantly upregulated in tumor cells (**Supplementary Table 7**).

### Multiple integrin genes upregulated in IO exposed cells in the stroma

Amongst stromal FOVs, *GPX3* is the most significant downregulated gene in IO exposed tumor cells (FDR <0.001), and *AZU1* was found to be downregulated in regulatory T cells (FDR < 0.001) and fibroblasts (FDR < 0.001; **Supplementary Table 8**). B cells showed the most genes (334 genes) expressed higher in IO exposed FOVs than IO naïve FOVs, while vasa recta endothelium, fibroblasts, tumor, and regulatory T cells also had high expression in many genes (158, 154, 130, and 123, respectively). Amongst stromal fibroblast cells *, ITGAV, ITGA1, ITGB1,* and *ITGB5* were significantly upregulated following IO and notably, isolated tumor cells found in the stroma showed upregulation of *B2M*, *VIM, ITGA5, ITGB2*, and *ITGA3* (**Supplementary Table 8)**.

The most significant changes in expression between sarcomatoid tumor FOVs and non-sarcomatoid tumor FOVs occurred in CD8 T cells, including downregulation of the anti-inflammatory RNA-binding protein *ZFP36* and downregulation of *TGFBR2* (**Supplementary Table 9**). Two genes (*EPOR* and *LIFR*) were significantly downregulated in stromal M2 macrophages in sarcomatoid tissues while no genes were significantly upregulated (**Supplementary Table 10**). B cells also showed eight genes upregulated in sarcomatoid samples, including *COL9A3* and *ITGB2* (FDR < 0.001 for both).

### Epithelial-mesenchymal transition (EMT) gene set spatially enriched in IO exposed samples

Tumor FOVs exhibited spatial enrichment of the EMT gene set, with spatial aggregation observed in 4 of 5 samples exposed to IO and only 1 of 6 IO naïve samples (**Table 2**). Expression of the hypoxia gene set was also spatially clustered in all TKI/IO exposed tissue samples; in contrast only 1 IO naïve sample showed spatial clustering. The hypoxia gene set was clustered in one patient who received ipilimumab/nivolumab (p = 0.001). Higher spatial clustering was also seen in IL2/STAT5 signaling (5 of 5 IO exposed p < 0.001; 1 of 6 IO naïve p < 0.1), IL6/JAK/STAT3 signaling (4 of 5 IO exposed p < 0.001; 1 of 6 IO naïve p < 0.1), MYC targets V1 (3 of 5 IO exposed p < 0.001; 2 of 6 IO naïve p ≤ 0.001), and oxidative phosphorylation amongst IO exposed samples (5 of 5 IO exposed p < 0.001; 0 of 6 IO naïve p < 0.1). Very few gene sets showed more clustering in IO naïve samples than samples exposed to IO (**Figure 3**).

**Figure 3:**
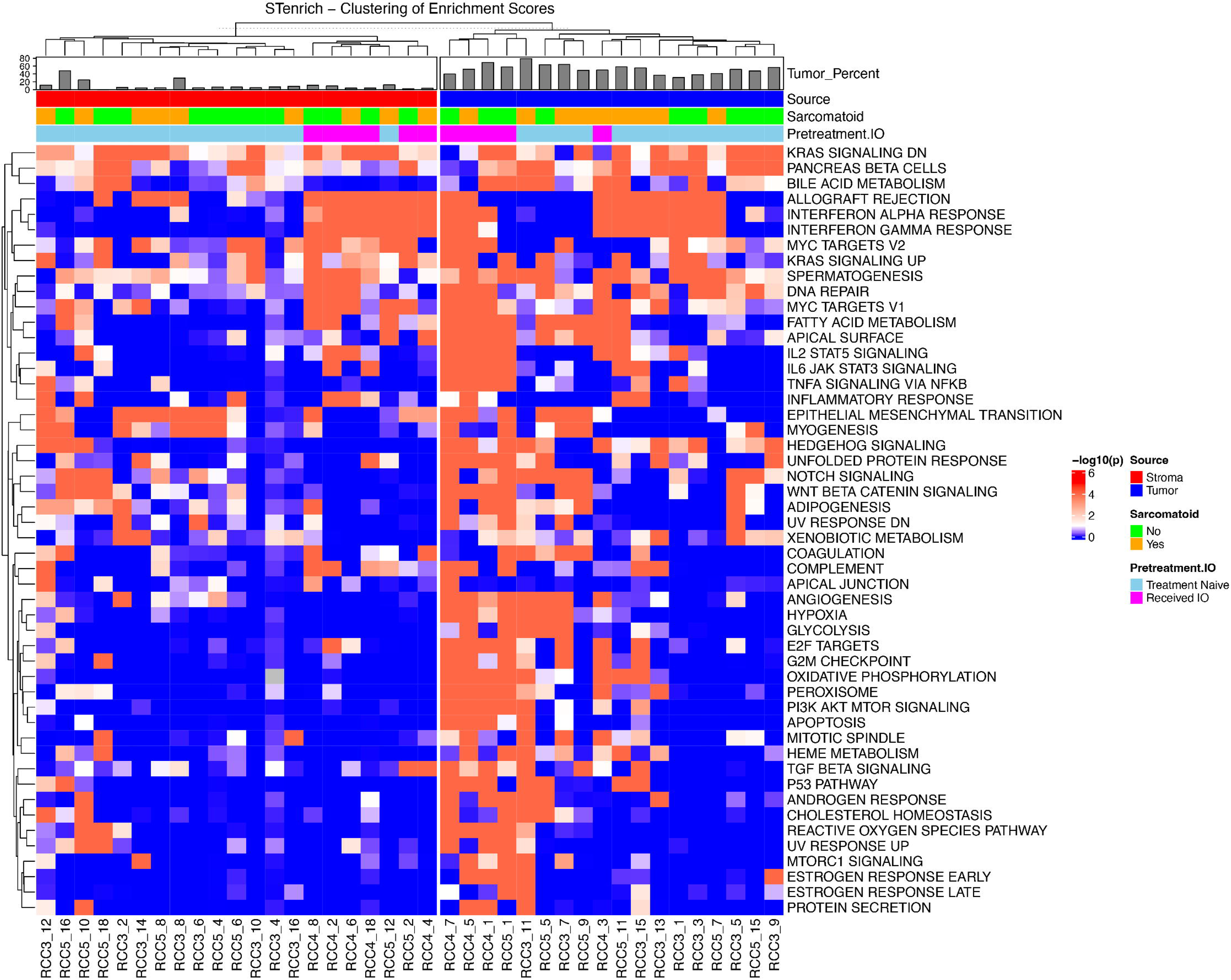
Spatial enrichment of cells with high (> mean + 1 standard deviation) Hallmark gene sets scores on tumor and stromal FOVs showing high spatial enrichment of gene sets in IO exposed tumor FOVs.

**Table 2:** Results of the spatial enrichment of Hallmark gene sets in all samples.

In stromal FOVs, interferon α and gamma response were clustered in all IO exposed samples with no clustering observed in IO naïve samples (**Table 2**). Both IO naïve and IO exposed groups harbored samples with cells that have high enrichment of EMT clustering. Genes related to IL6/JAK/STAT3 showed high spatial enrichment in 2 out of 6 IO exposed stromal samples and only in 1 of the IO naïve stromal samples.

In sarcomatoid tumor FOVs, TGF-β signaling was spatially enriched in 3 of 7 sarcomatoid samples, and none of the non-sarcomatoid samples. Genes related to the G2M checkpoint displayed significant spatial enrichment in 3 sarcomatoid samples and none of those without sarcomatoid features. Also, 4 sarcomatoid samples showed spatial enrichment of oxidative phosphorylation while no non-sarcomatoid showed spatial enrichment (**Table 2**).

In sarcomatoid stromal FOVs, TNFα signaling via NFKβ showed significant spatial enrichment in 3 of 7 sarcomatoid stromal samples and 0 of 8 non-sarcomatoid stroma samples along with the complement gene set (**Table 2**). Genes involved in adipogenesis demonstrated significant spatial enrichment in 5 of the 8 non-sarcomatoid samples and 2 of the 7 non-sarcomatoid (**Figure 3**).

### *COL4A1* and *ITGAV* spatially autocorrelated in the stroma of IO exposed samples

Moran’s I is a statistical measure that assess spatial autocorrelation between two variables, that is the degree to which high or low values of those two variables correlate across a Euclidean distance. A FOV in which two genes are highly expressed between many neighboring cells will generate an elevated Moran’s I (where values range from -1 to +1 with negative values indicating inverse spatial relationship and positive values indicating direct spatial relationship in gene expression). Moran’s I for ligand-receptor pair genes from the EMT pathway were not significantly different between IO naïve and exposed samples in tumor FOVs (**Supplementary Table 11**). The smallest FDR observed was between *TGFB1* and *ITGB5* (FDR = 0.91) where IO naïve samples averaged a slightly negative Moran’s I (I = -0.019) and IO exposed samples averaged slightly positive Moran’s I (I = 0.05). Ligand receptor genes in the IL6/JAK/STAT3 signaling gene set didn’t show strong differences in spatial autocorrelation between IO naïve and exposed samples (**Supplementary Table 12**). Level of autocorrelation between *TGFB1* and *ACVRL1* pair was the most different between IO naïve and exposed cohorts with FDR = 0.41.

Among the stromal FOVs, the *COL4A1* and *ITGAV* pair showed significant differences in Moran’s I between naïve and exposed samples (FDR = 0.056; **Supplementary Table 11**). IO naïve samples had an average Moran’s I = 0.0127 while those exposed to IO showed an average Moran’s I = 0.120, showing significantly higher spatial autocorrelation following exposure to IO treatment. *ITGAV* expression did not significantly differ between tumor cells in the stroma FOVs (FDR = 0.934; **Supplementary Tables 8, 11**).

In sarcomatoid samples, bivariate Moran’s I did not identify any ligand receptor pairs from either EMT or IL6 pathways that were spatially correlated in either the tumor or stroma FOVs (**Supplementary Table 11, 12**; **Supplemental Text)**.

### Fibroblasts, myofibroblasts, and endothelial cells displayed the highest ***COL4A1*** and ***ITGAV* expression**

*COL4A1* and *ITGAV* expression was plotted for the FOV with the greatest bivariate Moran’s I (RCC4 – FOV8) to identify cells associated with regions of high expression (**Figure 4A**). Myofibroblasts showed overlap with regions of high *COL4A1* expression and both myofibroblasts and tumor cells showed overlap with the areas of high *ITGAV* expression. Locations of all cells in the FOV is shown in **Figure 4B**. Boxplots were also constructed to link cellular phenotypes with greatest expression of *COL4A1* and *ITGAV*. This showed that fibroblasts, myofibroblasts, and kidney specific cell types (like tumor cells and endothelium) had the highest expression (**Supplementary Figure 2**). Tumor cells showed high expression of both *COL4A1* and *ITGAV*.

**Figure 4:**
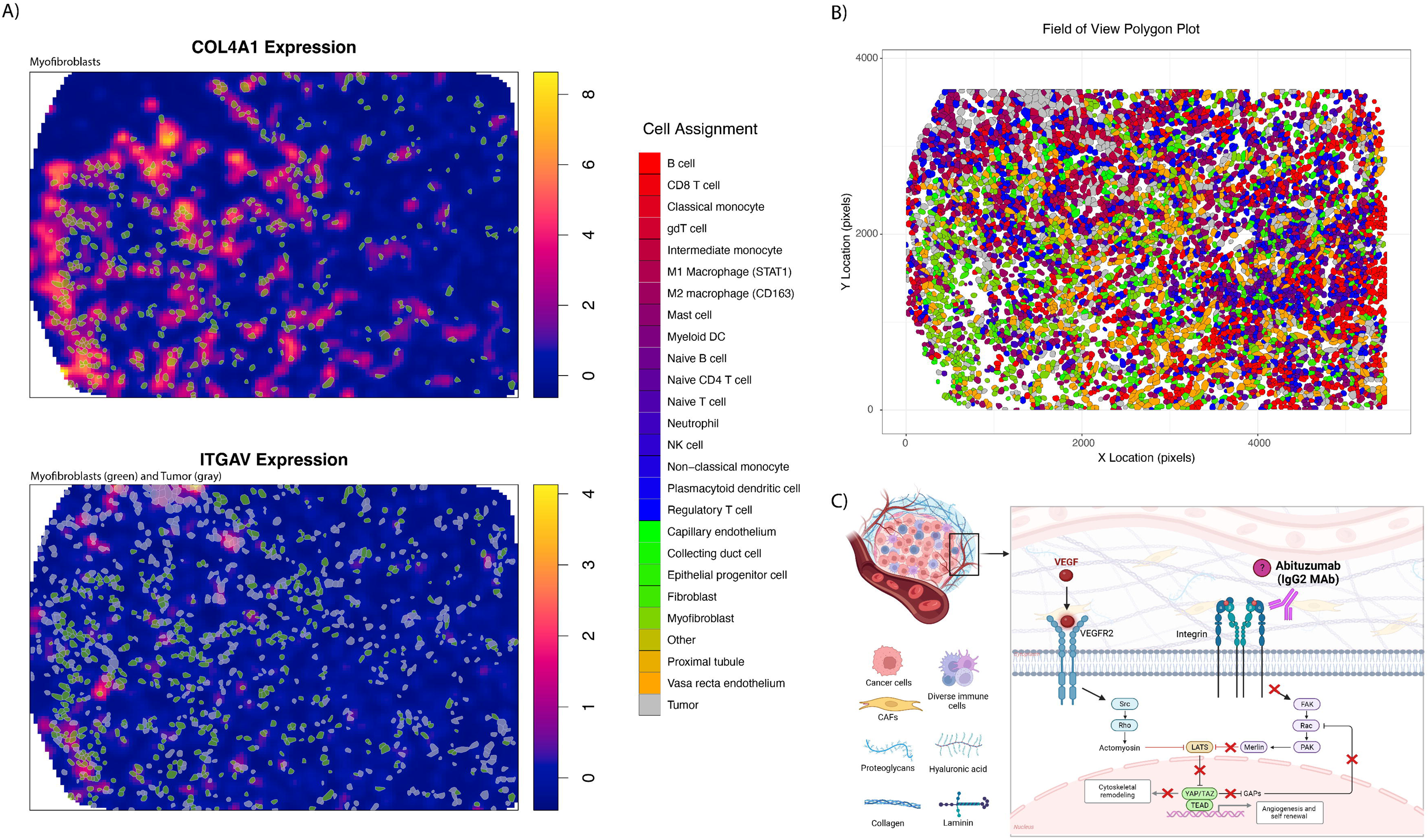
Expression of COL4A1 and ITGAV on RCC4 – FOV8. Figure A) shows overlap for myofibroblasts and tumor and high gene expression. Locations of all cells and their respective phenotypes can be seen in B). Pathway description of VEGF mediation neovascularization, of which integrin and YAP are involved.

### More fibroblasts express integrin αV in IO exposed samples on multiplex immunofluorescence (mIF)

To validate our findings of ligand-receptor gene expression in protein products, we conducted multiplex immunofluorescence on the spatial transcriptomics TMA cores. Abundances of integrin αV and type IV collagen α1 positive cells were compared between tissue groups (e.g., IO naïve vs IO exposed, sarcomatoid vs non-sarcomatoid) using beta-binomial modeling. We did not identify differences in the percent of cells that were integrin αV and type IV collagen α1 positive on tumor FOVs between IO naïve and IO exposed nor between primary IO naïve and sarcomatoid samples. However, as observed with the IO naïve vs IO exposed bivariate Moran’s I analysis using gene expression, integrin αV positive cells on mIF were significantly more abundant after exposure to IO in the stroma as compared to IO naïve tissues (p = 0.001; **Supplementary Table 13**).

Next, we set out to determine which cell type was contributing to the difference of integrin subunit αV protein in the IO exposed samples. IO naïve and IO exposed samples showed a significant difference in fibroblast (SMA marker) cells that are integrin αV positive in the stromal compartment (p < 0.001; **Supplementary Table 14**). A similar trend was identified in the tumor compartment (p = 0.064).

Compared to IO naïve samples, the proportion of the area positively stained for type IV collagen α1 (including extracellular protein) was higher amongst IO exposed samples in both tumor and stromal regions, but this was not statistically significant (p = 0.9 and p = 0.4).

### Type IV collagen expression higher in advanced ccRCC stage; COL4A1 and ITGAV have the highest expression in kidney cancer compared to other cancers

To further characterize the presence of *COL4A1* and *ITGAV* in ccRCC we examined the Clinical Proteomic Tumor Analysis Consortium (CPTAC), The Cancer Genome Atlas (TCGA) and the Genotype-Tissue Expression (GTEx) project. ^19–21^ Examining normal tissue samples from GTEx, we found kidney and fibroblast cells demonstrated some of the highest expression of these genes compared to other cell types (**Supplementary Figure 3**). In CPTAC, type IV collagen protein expression was significantly higher in advanced ccRCC disease (stage IV) compared to localized disease (stages I, II, III vs IV; p = 0.015; **Figure 5**).

**Figure 5:**
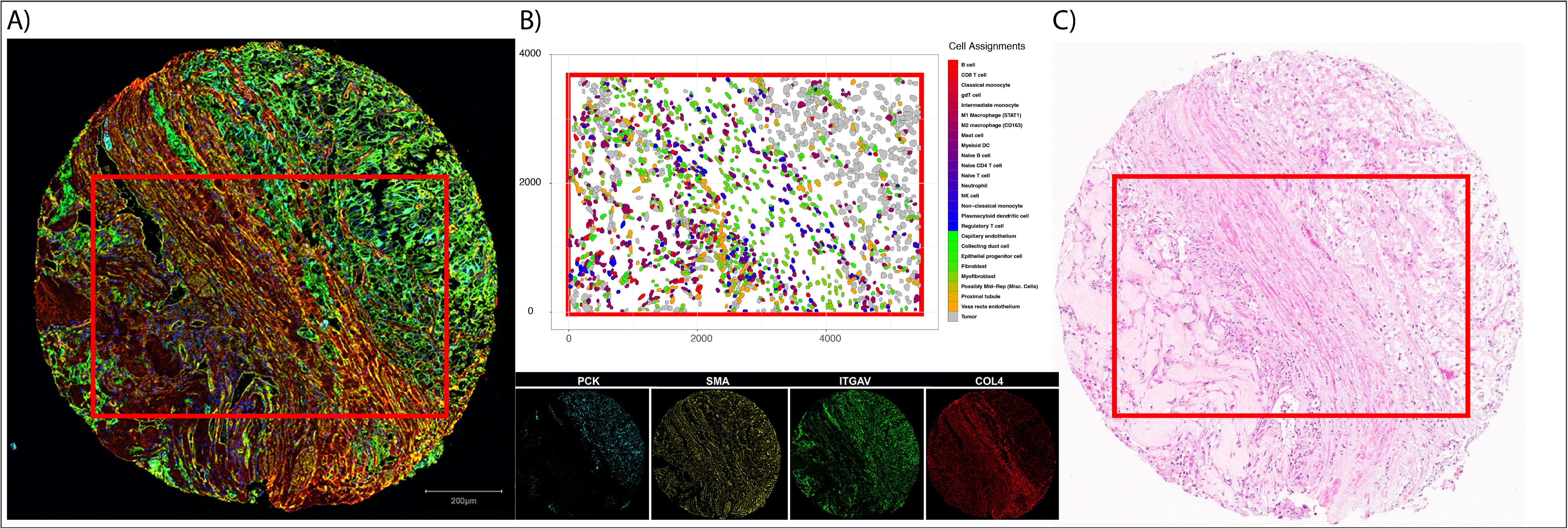
Example core that subjected to different assays. A) displays multiplex immunofluorescence staining for pancytokeratin (endothelium, PCK), smooth muscle actin (fibroblasts, SMA), integrin subunit alpha (ITGAV), and type 4 collagen (COL4). Cell types derived from CosMx SMI gene expression (B) show structure identified in both the multiplex immunofluorescence image and H&E (C).

**Figure 6:**
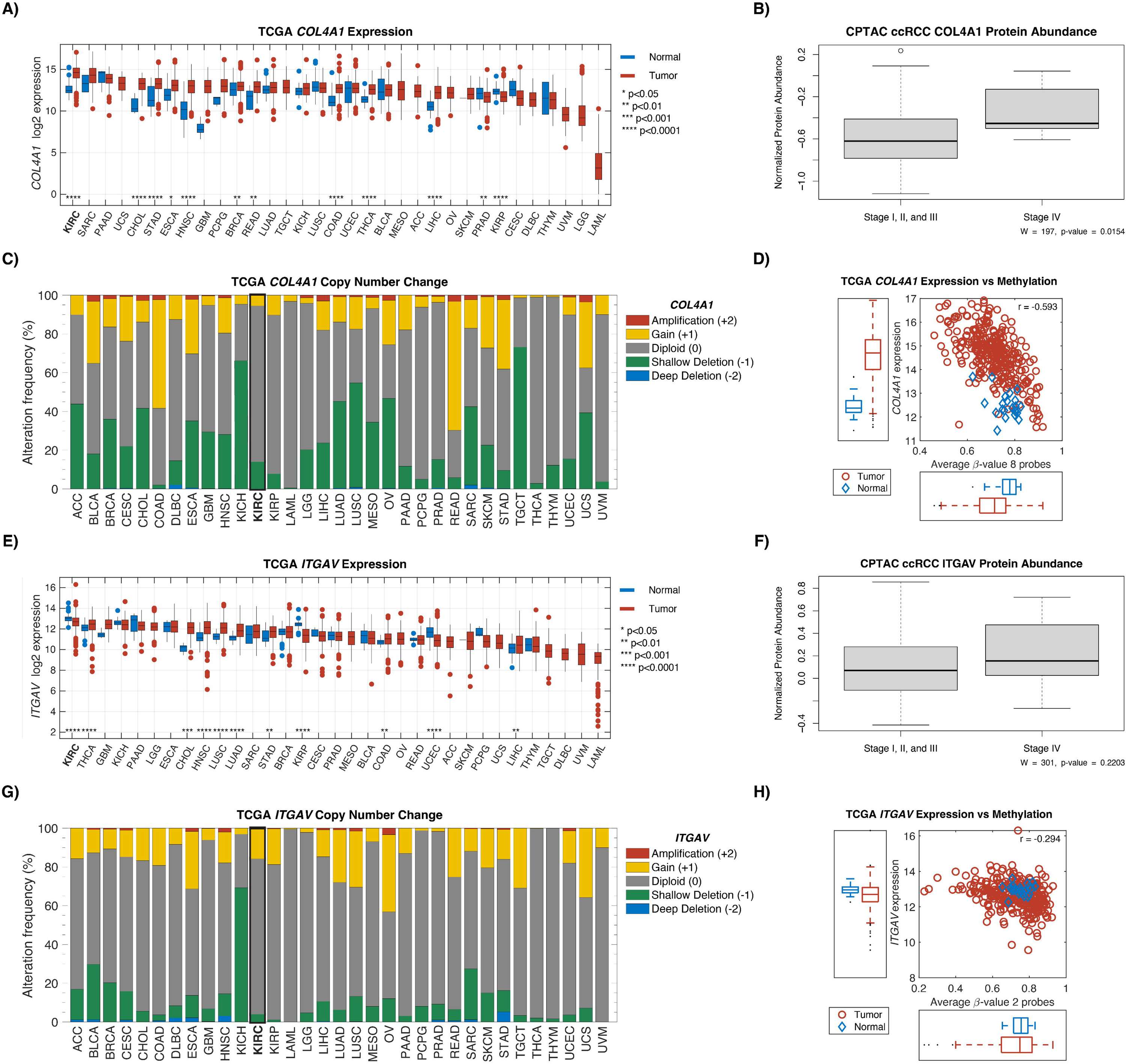
Exploration of COL4A1 and ITGAV in TCGA and CPTAC. Gene expression in tumor and normal for COL4A1 (A) and ITGAV (E). Protein abundance between low stage ccRCC (I, II, and III) and high stage ccRCC (IV) for COL4A1 (B) and ITGAV (F). C) and G) show the copy number change associated with the two genes. TCGA gene expression against methylation levels are shown in D) and H).

Using the PanCan Atlas from TCGA, expression of ITGAV and COL4A1 was found to be the highest in ccRCC specifically, compared to all other cancers. The majority of copy number variations (CNVs) for these genes were found to be diploid. Interestingly, *COL4A1* gene was found to be significantly hypomethylated in ccRCC tumor samples compared to adjacent normal tissue (**Figure 5**).

## Discussion

Given the lack of clinically validated biomarkers in the prediction of ccRCC IO efficacy, the cell diversity and spatial heterogeneity of the TIME holds much promise in the extraction of clinically meaningful information from biopsy or surgical tumor samples. By leveraging differences between treatment naïve and eco-evolutionarily selected residual tumors following IO therapy, we sought to reveal unique characteristics of the IO resistant TIME using cellular resolution spatial transcriptomics. Most cell abundance and gene expression changes occur within the stroma compartment of the tumor tissue amongst immune cells. Spatial gene set enrichment analysis at cellular resolution revealed enrichment of EMT and IL6 hallmark gene set pathways. Uniquely, we identified an increase in the stromal colocalization of *ITGAV* and *COL4A1* transcripts moving from the IO naïve to IO exposed setting which represents a unique spatial relationship not previously identified in ccRCC. Finally, we validated these associations at the protein level through multiplexed immunofluorescence.

High EMT enrichment scores on bulk RNA sequencing are associated with worse disease specific survival in ccRCC ^22^, and various EMT ligand-receptor pairs have been implicated.^23, 24,25^ Prior spatial transcriptomic work using spot-based resolution has also identified EMT-rich tumor cells being associated with worse prognosis in the TCGA cohort. ^26^ Still, specific ligand-receptor pairs from the EMT pathway were not elucidated in these studies. This may reflect the limitation of scRNA-seq data or, in the case of spot-based spatial transcriptomics, the loss of cell-to-cell spatial information. Although average ligand expression is high in one cell type and receptor expression high in another, this does not demonstrate colocalization and the likely affinity for biologic cross talk. In this study, we describe a GSEA method that also incorporates spatial data. We hypothesize that the significance of spatial gene set enrichment represents downstream activation of cell signaling molecules. Within specific gene sets, we were able to identify a curated list of known ligand-receptor pairs and identify one such signaling pair which was also spatially enriched and unique to IO exposed tumors.

EMT activation may confer resistance to therapy in cancer cells by a number of biological pathways, including cell cycle arrest, alteration of cellular transporters, and dampening the cytolytic activities of CD8+ T cells. ^27^ *YES1*, the most significantly upregulated gene amongst IO exposed tumor cells in this cohort, may explain one specific mechanism via YAP1 (YES-associated protein), which is also associated with poor survival in treatment naïve ccRCC patients.^28, 29^ In human ccRCC cell lines, Chen et al. demonstrated EMT enrichment in cancer cells with upregulated *YAP1*.^30^ Given our findings of the upregulation of *YES1* in IO exposed tumor cells, this remains a plausible mechanism in our cohort. Moreover, the Hippo-YAP pathway is a known linchpin in therapy resistance for many cancers and multiple major drug classes, including immunotherapy.^31^ YAP1-mediated immune-resistance is possibly conferred by increased expression of PDL1 in other cancers. *CD274* expression (gene for PD-L1) was not significantly higher amongst tumor cells in this study, however, PD-L1 has also not been shown to be a reliable biomarker for immunotherapy response in ccRCC. ^32^ Thus, additional studies of *YAP1* and related Hippo signaling pathways are needed in this population using targeted gene panels.

Beyond *YES/YAP1*, EMT has been shown to more directly confer IO resistance by other mechanisms. In breast cancer models, where epithelial tumor cells were infiltrated by CD8+ T cells, mesenchymal tumors contained regulatory T cells and M2 macrophages and were resistant to IO treatment with checkpoint blockade. ^33^ Additionally, mixed tumors containing only a fraction of mesenchymal cells continue to recruit regulatory T cells and M2 macrophages to the primary tumor, in line with the concept of immune cell exclusion.^33, 34^ Inferred and predicted ligand-receptor pairs have been suggested but none with cell-to-cell spatial resolution.

We demonstrated that stromal colocalization of *ITGAV* and *COL4A1* in the IO exposed TIME is increased among fibroblasts, tumor cells, and endothelial cells. On mIF validation testing, *ITGAV*+, *SMA*+*ITGAV*+ in the stroma were also seen in higher abundance following IO, suggesting a proteomic correlative increase in integrin αV amongst stromal fibroblasts. Further validation testing from the CPTAC dataset indicated collagen IV α1 protein was enriched in advanced stages of disease (**Figure 5B**). Methylation data from TCGA data suggests this enrichment may be a result of DNA hypomethylation (**Figure 5D**).

The *ITGAV* protein (integrin subunit αV) has a reported role in cell migration and metastasis in several cancers.^35–37^ In ccRCC, integrin overexpression has been associated with tumor grade, distant metastases, and overall survival (OS). ^38–42^ It has also been described in mTOR inhibitor resistant tumors analyzed with flow cytometry.^43^ The role of *ITGAV* specifically in ccRCC has not yet been well characterized, however, Crona et al. identified an intronic variant of *ITGAV* that leads to overexpression and associated with decreased OS in TKI treated patients. ^42^ Feldkoren et al. used human renal cell carcinoma cell lines to show TGF-β dependent overexpression of the integrin αv-β3 leads to decreased E-cadherin expression and increased cell mobility.^41^ Although the biological basis of *ITGAV’s* contribution to ccRCC tumorigenesis has not yet been fully elucidated, experiments in other cancers have demonstrated positive association with well described ccRCC pathways, such as VEGF mediated neovascularization (**Figure 4C**).^44–48^ Inhibition of *ITGAV* also appears to increase T cell killing of melanoma cells in vitro. ^49^ Thus, the current study adds to an existing biological rationale for investigating therapeutic targets of this signaling pathway, especially among patients with ccRCC treated with IO.

Integrin targeting has been explored in other cancers. ^36^ Multiple agents have been developed against the integrin αv subunit, including cilengitide and abituzumab, and studied in phase II trials.^36, 50, 51^ Oncologic outcomes with integrin targeting in other settings have been underwhelming thus far. In the only phase III study in glioblastoma, combination therapy with cilengitide did not improve survival over standard of care. ^50^ Even so, integrin targeted therapeutics have not been tested in the IO exposed setting in kidney cancer. This represents a potential area of combination or sequential treatment with already developed therapeutics. More studies are needed in this area.

The presence of extracellular matrix (ECM) components such as collagen IV and cancer associated fibroblasts (CAFs) have been found to play an integral role in tumor growth, migration, and neovascularization in ccRCC.^52^ ECM-rich gene signatures have been shown to be associated with poorer overall survival. ^53^ Collagen IV specifically is a known VHL interaction partner, and found in abundance in ccRCC.^54^ Yet the majority of ECM is produced not by cancer cells but other stromal cell types such as fibroblasts. This fibrotic overproduction of collagen and other ECM proteins appears to be present in multiple kidney-specific pathologies, as validated in recent spot based spatial transcriptomics. ^55^ In cancer, CAFs have been spatially associated with mesenchymal like cells in ccRCC at the tumor-stromal interface on both spot-based spatial transcriptomic and proteomic analysis. ^56^ Thus, our findings validate this specific interaction with cellular resolution and highlight the increased autocorrelation in the IO resistant TIME.

Our analysis also identified an increased number of dysfunctional CD8+ T cells within the stroma of the IO exposed ccRCC TIME. ^57–59^ The shift of functional to dysfunctional CD8+ T cell infiltration within the TIME over pseudotime has been well characterized.^6, 57, 60–62^ The upregulation of T cell exhaustion markers (*TOX*, *EOMES*), as well as functional T cell responsiveness (*GZMK*) in the IO-resistant TIME in our analysis supports prior findings. ^60^ The mix of exhaustion and functional markers representing both exhaustion and responsiveness may reflect the unique clinical scenario of the patients from which these samples are derived, somewhere in between complete response and secondary refractive disease.

Among sarcomatoid samples, we found a higher abundance of M2 macrophages in the stroma, which is consistent with prior studies. ^63^ The most significant gene expression differences in tumor FOVs are found in CD8+ T cells, M2 macrophages, and regulatory T cells. Interestingly, *TGFBR2*, was downregulated in tumor embedded CD8 T cells. This gene has been shown to be associated with T cell exhaustion in breast cancer. ^64^ Thus, its downregulation in sarcomatoid tumors might be one plausible mechanism for sarcomatoid tumors having favorable responses to immunotherapy. TGF-beta signaling in T cells is known to regulate another significantly downregulated gene in this study, *ZFP36*, which encodes the anti-inflammatory RNA-binding protein tristetraprolin.^65^ Tristetraprolin is suppressed in multiple aggressive cancers, such as breast and prostate, and can be associated with inflammatory subtypes.^66^ Its dysregulation in T cells specifically is often seen in inflammatory conditions, such as rheumatoid arthritis, and multiple sclerosis.^67^ However, the role of tristetraprolin in cancer-associated CD8 T cells is still unclear. Additional study is required to illuminate how this specific CD 8 T cell profile might confer an exceptional response to IO therapy.

The overall role of TGF-β signaling in sarcomatoid tumors has not been conclusive.^68^ Traditional GSEA of bulk RNA sequencing have not shown enrichment in the TGF-β gene set, however, Wang et al. observed upregulation of the TGF-β signaling pathway across multiple sarcomatoid samples using commercial network-based pathway analysis. ^63, 69^ The contrasting results of these two studies may be rooted in their differing methods. In the current study, we found that 3 out of 6 sarcomatoid samples were spatially enriched for TGF-β signaling, while none of the non-sarcomatoid samples were enriched. Yet in differential gene expression analysis by cell phenotype, neither TGF-β1 nor its receptor was significantly changed in expression in sarcomatoid samples. These conflicting results may represent the limitation of pathway analysis all together or a limitation of use of partial gene sets. Although the promise of spatial transcriptomics lies in its high-plex capacity, only a portion of the hallmark gene sets could be analyzed in this current study. Additional studies, likely with expanded gene targets, are needed to evaluate more components of the TGF-β signaling pathway and their biologic role in sarcomatoid disease.

Limitations of this study include the relatively small number of patient samples and a mixed population of treated patients including TKI exposed patients. Sample size limitations, however, is bolstered by the single cell level analysis of multiple tissue regions and the use of contemporary treatment regimens. Several bioinformatics steps were developed in this study, including malignant cell phenotyping and spatial GSEA, and require validation in external datasets. Additionally, transcriptomic analysis was limited to genes present on the CosMx SMI platform.

## Conclusions

Using cellular resolution spatial transcriptomics in ccRCC, we found that IO exposure is associated with increased spatial gene set enrichment of the EMT pathway and colocalization of ligand-receptor transcripts *COL4A1* and *ITGAV*. The cell types with the highest expression of these two genes were fibroblasts, tumor cells, and other endothelium cell types which was also seen on multiple immunofluorescence. Additional study is needed to elucidate the biological basis for this shift in the ccRCC TIME, as well as examining the possible therapeutic potential of integrin following IO treatment.

## Methods

### Patient Cohorts

We prospectively collected tumor samples from 21 patients with ccRCC (MCC #20148, Advarra (Pro00038234)). The presence of sarcomatoid elements was identified on nine patients (2x IO exposed and 7x IO naïve) and 12 patients without (4x IO exposed and 8x IO naïve). Among the six patients who had tumors collected after IO-based therapy: four patients received the combination of pembrolizumab and axitinib for a median time on therapy of 8.5 months and two patients received nivolumab and ipilimumab for a median of 5.5 months. These patients selected for surgical resection of their primary tumor due to concerns for remaining viable disease after initiation of IO therapy. Eight tumor samples were collected from patients without sarcomatoid elements and from those who presented with localized disease initially and underwent surgical resection (i.e., IO naïve tumors). Clinical attributes of all patient samples can be found in **Table 1**. A trained pathologist (JD) identified final paired tumor and stroma (tumor-adjacent) samples: 8x IO naïve non-sarcomatoid, 6x IO exposed, and 7x IO naïve sarcomatoid.

### Tissue microarray (TMA) construction and CosMx SMI spatial gene expression profiling

Each patient’s tumor sample had two spatially distinct tumor and stroma tissue samples prepared using 1mm core biopsies for TMA creation. A total of 42 core formalin-fixed paraffin-embedded (FFPE) samples from 21 patients allocated across 3 slides for TMA creation. We utilized a protocol allowing for 20 field of views (FOVs) at 0.9mm x 0.7mm that were profiled with the scanning area per slide. Full details of the CosMx SMI chemistry and workflow can be found in He et al 2021. ^70^ Briefly, each slide underwent in-situ hybridization (ISH) of 978 mRNA probes optimized to investigate the biology of single cells across tumors and diverse organs. Of these probes, 758 genes were selected to capture critical cell states and cell-cell interactions. The remaining genes and markers were selected to optimize the panel’s power to distinguish between different cell types. A list of probe targets can be found in **Supplemental Table 15**. The four markers used were CD3 (T cells), CD45 (lymphocytes), CD298 (ubiquitous human cell membrane protein), and pancytokeratin (epithelial cells) for multiplex immunofluorescence. These markers were also used for multimodal cell segmentation which was provided by Nanostring.

### Data Quality Control and Non-Tumor Cell Phenotyping

For each tissue sample, probe counts, spatial coordinates, and cell segmentation output was obtained and processed using open-source packages implemented in R v4.3.0 (*Seurat* v4.3.0, *InSituType* v1.0.0, and *spatialTIME* v1.3.3.3).^17, 71, 72^ Cells with less than 20 transcript counts or abnormally large cell area (resulting from segmentation errors and defined as greater than 5 times the geomentric mean) were removed from downstream analyses. ^73^ Additionally, FOVs with less than five cells segmented were considered as failed assays and removed from data. Counts were normalized with ‘SCTransform’ allowing removal of technical cell-to-cell variation beyond simple log-transformations.^74^ Using the transformed expression data and the 4 protein fluorescence data, the *InSituType* software was used to phenotype cells based on the Kidney Cell Atlas Single Cell reference. ^75^ Prior to analysis with InsituType, normalized gene expression of all cells was scaled to the largest library size (i.e., cell with highest gene counts), followed by averaging of gene expression in the Kidney Cell Atlas by cell phenotype.

Following *InSituType* phenotyping, T cells and mononuclear phagocyte (MNP) clusters were subset and identification of sub-clusters was performed on that subset with principal component analysis (PCA) and UMAP projections. Differentially expressed genes (DEG) among sub-cluasters were detected with the “FindAllMarkers” function within *Seurat*. The DEG were used to further refine phenotypes (i.e., regulatory T cells, Naïve CD4+ T cells). Marker genes with positive log fold change (LFC) were queried in the Human Protein Atlas single cell expression data to assign cluster identity.

### Identification of Malignant Cells

Given that our kidney cell reference atlas only includes healthy cells, we could not use it for accurate identification of malignant cells. To identify malignant cells (tumor) we implemented an approach based on lasso logistic regression of tumor marker expression. Since VEGFA expression has been shown to be upregulated in patients with ccRCC ^76^, we first selected a subset of cells from stroma and tumor FOVs with distinct proximal tubule expression profiles (i.e., high or low *VEGFA* expression; **Supplementary Table 3**). Differential gene expression analysis was performed between tumor and stroma proximal tubule cells (cell of origin for ccRCC) to determine marker genes. Proximal tubule cell expression and tumor/stroma assignment based on *VEGFA* expression was passed to the function ‘cv.glmnet’ from the R package *glmnet* (v4.1-7) to train a lasso logistic regression model with 10 fold cross validation (L1 regularization) and providing a stronger, smaller marker gene list. ^77^ Using the lasso determined marker genes, a generalized linear model (GLM) was fit and a threshold of 0.5 was used to classify a cell as tumor (< 0.5) or non-tumor (>= 0.5). This model was then applied to all cells where the phenotypes were determined from *InSituType* to be kidney tissue related (i.e., cells determined not to be fibroblast/myofibroblast or immune-related). Additionally, on stromal FOVs where the cell assignment was ‘glomerular endothelium’ from *InSituType* and the lasso model classified as ‘tumor’ cell, we reassigned these cells as glomerular endothelium upon review with a genitourinary pathologist. Lastly, cell types from the Kidney Cell Atlas were collapsed to increase number of cells in each cell type (**Supplementary Table 16**). *InSituType* phenotype assignment for glomerular endothelium were used on stromal FOVs when our GLM predicted cells to be malignant follow consultation with a pathologist.

### Differential Cell Type Abundances

To explore differences in cell abundance between the patient groups, beta binomial models were used to test for relationships between the number of positive cells and the treatment for primary disease, as well as IO naïve sarcomatoid classes. P-values were adjusted with the Benjamini-Hochberg ^78^ false discovery rate (FDR) and a threshold of 0.1 was used to identify statistical significance.

### Differential Univariate Clustering of Phenotypes

The level of spatial aggregation among cells of a given type was measured to determine differences associated with IO exposure or presence of sarcomatoid features. To do this, we implemented Ripley’s *K* count statistic form the *spatialTIME* package in R for each cell type at a radius of *r* = 150 (27um), and adjusting for core-specific complete spatial randomness measurement (Ripley’s *K* measured across all cells in the FOV).^79–81^ The resulting measure is the “Degree of Clustering Exact” (DOCE) and allows for comparison of values between tissue samples by removing bias introduced by areas on FOVs where stationarity of the point process is violated (often due to cells not being measured in those areas). Missing measurements of cells may be due to squished tissue, FOV region expanding outside the tissue core, or necrotic tissue. The DOCE values were compared between the patient groups using a linear model predicting the DOCE from each patient group.

### Differential Gene Expression

We collapsed gene expression down to the FOV level by averaging all cells within a FOV (‘pseudo-bulk’) and conducting two-sample T tests between sample groups. ^18^ Additionally, we performed differential gene expression at the cell-level with ‘FindAllMarkers’ to see if genes were differentially expressed between sample groups (e.g., *MS4A1* in B cells). To perform this analysis, a linear mixed effects model was used whereby a random-effect was used to account for cells coming from the same FOVs (*lmerTest* v3.1-3).^82^

### Spatial Relationship of Enrichment Scores

To identify gene sets showing spatially aggregated (i.e. “hotspots”) enrichment scores in tumors from the three patient cohorts, we implemented a modified version of ‘STenrich’ from the *spatialGE* R package (v1.0.0). ^83^ Briefly, at the FOV level, cells with enrichment scores greater than 1 standard deviation above the mean were identified and the Euclidean distance between these identified cells was summed. The same number of identified cells were then randomly permutated 1000 times and the Euclidean distance between these permutated cell locations were summed to create an empirical null distribution. This empirical distribution representing the null hypothesis of no spatial aggregation was used to determine empirical p-values for the alternative hypothesis of spatial aggregation. The spatial enrichment p-values were then compared between patient cohorts by counting number of samples that showed significant evidence of enrichment hotspots.

### Identification of Ligand-Receptor Spatial Autocorrelation

Spatial proximity of ligand-receptor genes was evaluated using bivariate Moran’s I. We queried genes belonging to the epithelial mesenchymal transition (EMT) and IL6-JAK-STAT3 signaling pathways in the ligand-receptor pairs from *CellTalkDB*.^84^ For each cell, the three nearest neighbors were identified with the *spdep* R package (v1.2-8) (binary weigh = 1). ^85^ The selection of a low number of nearest neighbors was used to ensure the association of receptor expression considered only cells in close proximity to the ligand cell. To calculate the bivariate Moran’s I, the ligand and receptor gene expression and weights were input to the ‘moran_bv’ function in *spdep*. Bivariate Moran’s I ranges from -1 (strong inverse spatial autocorrelation) to +1 (strong positive spatial autocorrelation), with values around 0 indicating no spatial relationship between the gene expression values. To test for significant differences in Moran’s I value between the patient cohorts, two sample T-tests were performed.

### Validation of Ligand-Receptor with Multiplex Immunofluorescence

To evaluate the protein expression of spatially autocorrelated transcripts, we performed multiplex immunofluorescence. Tissue samples obtained from the same TMA cores used for SMI underwent mIF using a previously described protocol. ^86^ In brief, FOVs were stained for a panel of antibodies against PCK, smooth-muscle actin (SMA), *ITGAV* protein (integrin αv subunit), and *COL4A1* protein (collagen IV), as well as DAPI nuclear counterstain. Cell positivity for specific markers was set based on previously published staining patterns and visual intensities.^87^ We tested for differences in abundance of cells positive for *ITGAV* protein between patient cohorts, using non-parametric Wilcoxon Rank Sum tests (IO naïve vs IO exposed and IO naïve vs sarcomatoid). Due to elevated expression of *COL4A1* and *ITGAV* gene expression with tumor and myofibroblasts/fibroblasts in the bivariate Moran’s I analysis, we looked for associations with protein expression of *ITGAV* on these cell types (myofibroblasts and fibroblasts identified with antibody against SMA; tumor cells identified with antibodies against PCK; **Key Resources**). These associations were measured using beta-binomial models to look at positive cells between the patient cohorts with the *VGAM* R package (v1.1-8). ^88^ The *COL4A1* protein is largely extracellular, hence the total area proportion positive staining area determined visually across multiple samples was compared between IO naïve and IO exposed groups using Wilcoxon Rank Sum tests.

### CPTAC validation

Pre-processed, normalized, protein-level proteomic and phosphoproteomic data from the Clinical Proteomic Tumor Analysis Consortium (CPTAC) clear cell renal cell carcinoma (ccRCC) study ^19^ for 110 patients was obtained from www.linkedomics.org ^89^ on 7/31/2023. We compared COL4A1 and ITGAV protein abundances in tumors across stages I-IV using non-parametric Wilcoxon rank sum tests.

### Pan-Cancer Analysis and TCGA validation

All TCGA Pan-Cancer data was all downloaded from the National Cancer Institute’s Genomic Data Commons PanCan Atlas.^20^ Tumor type and sample type were inferred from TCGA barcode. The TCGA RNAseq data was extracted and log2 transformed. The DNA copy number variation (CNV) data (GISTIC data) were also extracted. Using the GISTIC2.0 DNA copy number analysis, the following numeric values were assigned with the cBioPortal interpretation presented in parenthesis: −2 = Deep Deletion (possibly a homozygous deletion); −1 = Shallow Deletion (possible a heterozygous deletion); 0 = Diploid; +1 = Gain (a few additional copies, often broad); +2 = Amplification (more copies, often focal). For data present in this chapter, only samples with Illumina’s Infinium HumanMethylation450 BeadChip data were used. Raw IDAT files were downloaded from TCGA. Preprocessing the data included normalization via internal controls followed by background subtraction using the methylumi R package.

**For a full list of resources, please see Key Resources table.**

### Declarations

Ethics approval and consent to participate:

Consent to participate was acquired through IRB under the Total Cancer care protocol (MCC #20148, Advarra (Pro00038234)).

### Consent for publication

Not applicable

### Availability of data and materials

Data produced by Nanostring CosMx SMI platform is available from BJM upon request and made public on date of publication, or earlier by request of reviewers. Original code for reproducing the analysis of this study can be found at: https://github.com/FridleyLab/ccRCC_CosMx_SMI_2023

The repository name is listed in the key resources table. Additional information required to reanalyze the data reported is available from BJM upon request.

### Funding

2023 Moffitt Team Science-Miles for Moffitt Award: P30 CA 076292

### Authors’ Contributions

Conception of study: AS, NHC, AEB, PAS, JD, BLF, and BJM. Tissue analysis: JN, CM-S, NLF, JAB, YCP, JD, WG, NS, SK, MG. Data Analysis: AS, MTH, OO, AEB, PAS, JD, BLF, and BJM. Manuscript construction: AS, MTH, TCP, OO, NHC, AEB, PAS, JC, RL, KYT, JM, BLF, and BJM. There were no non-author contributors. BJM accepts full responsibility for the work and/or the conduct of the study, had access to the data, and controlled the decision to publish.

## Supporting information

Key Resources

## Acknowledgements

This publication is supported by Tissue Core, Biostatistics and Bioinformatics Shared Resources, the 2023 Moffitt Team Science-Miles for Moffitt Award, and the Cancer Center Support Grant at the H. Lee Moffitt Cancer Center and Research Institute, an NCI designated Comprehensive Cancer Center (P30 CA 076292). We would also like to acknowledge Nanostring for providing technical support.

---

**Supplementary Figure 1:**
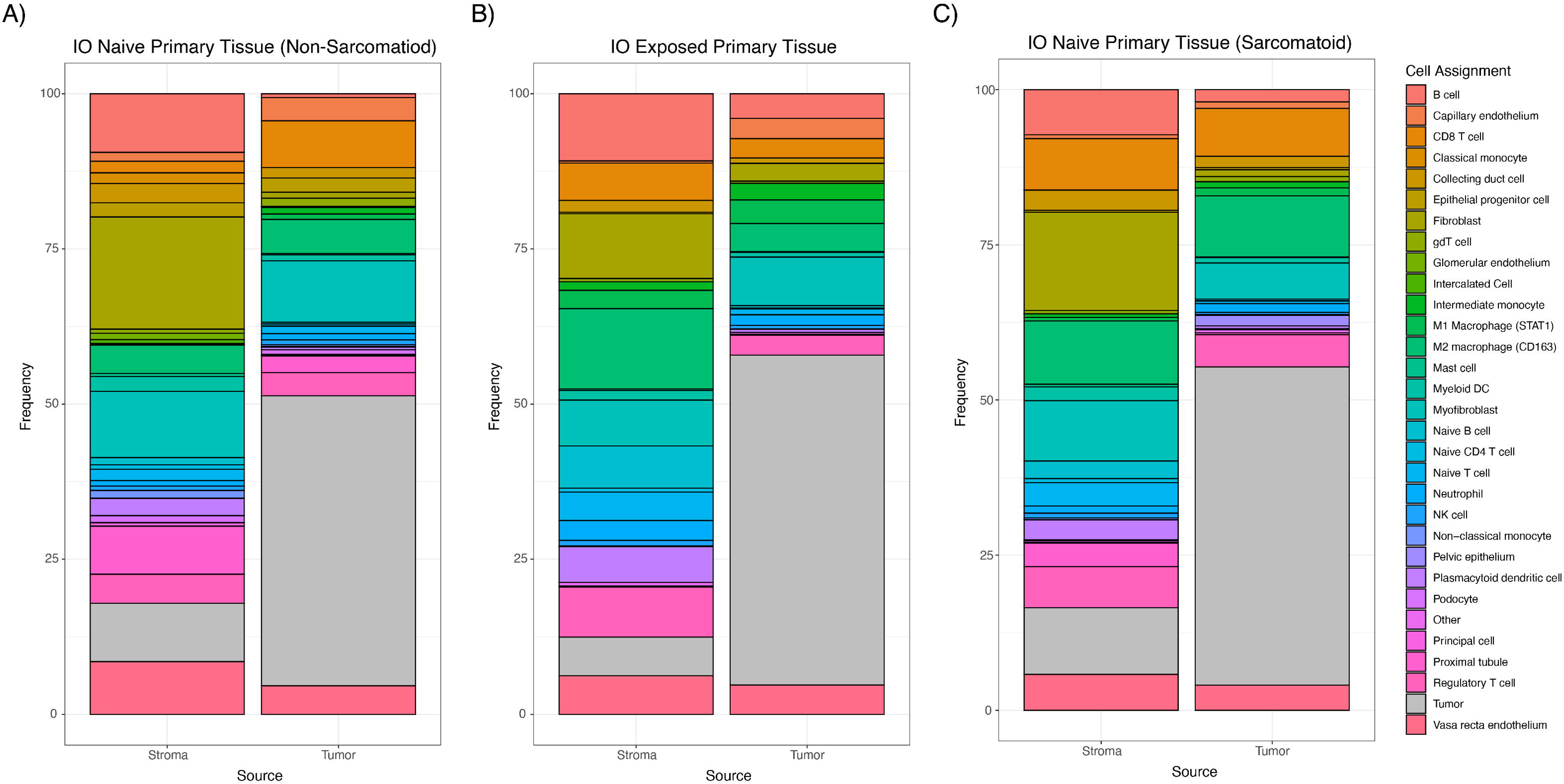
Abundance of final cell assignments. A) shows IO naïve primary non-sarcomatoid, B) shows IO exposed, and C) shows IO naïve primary sarcomatoid stroma and tumor FOVs.

**Supplementary Figure 2:**
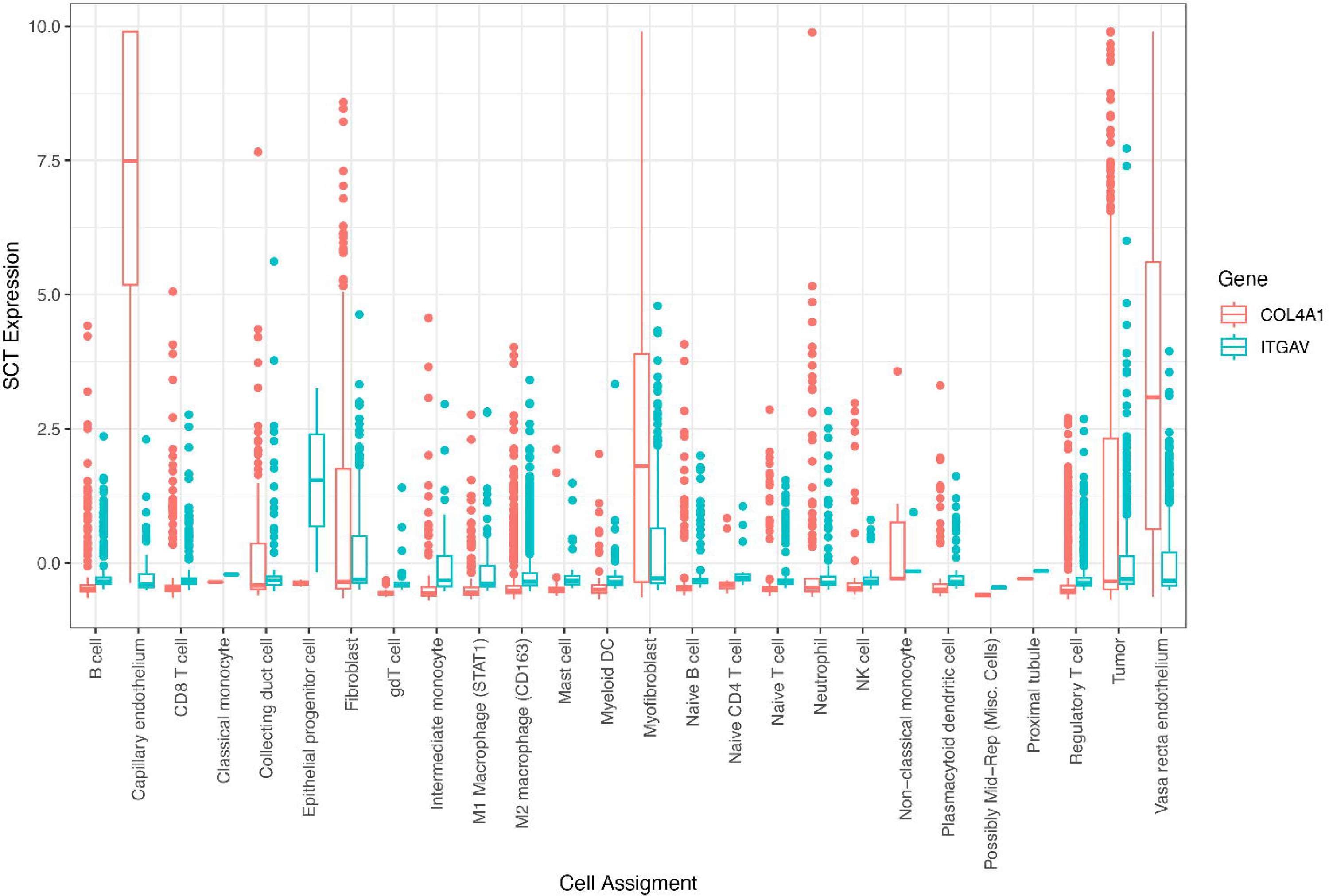
Expression of COL4A1 and ITGAV for cell types on RCC4 – FOV8.

**Supplementary Figure 3:**
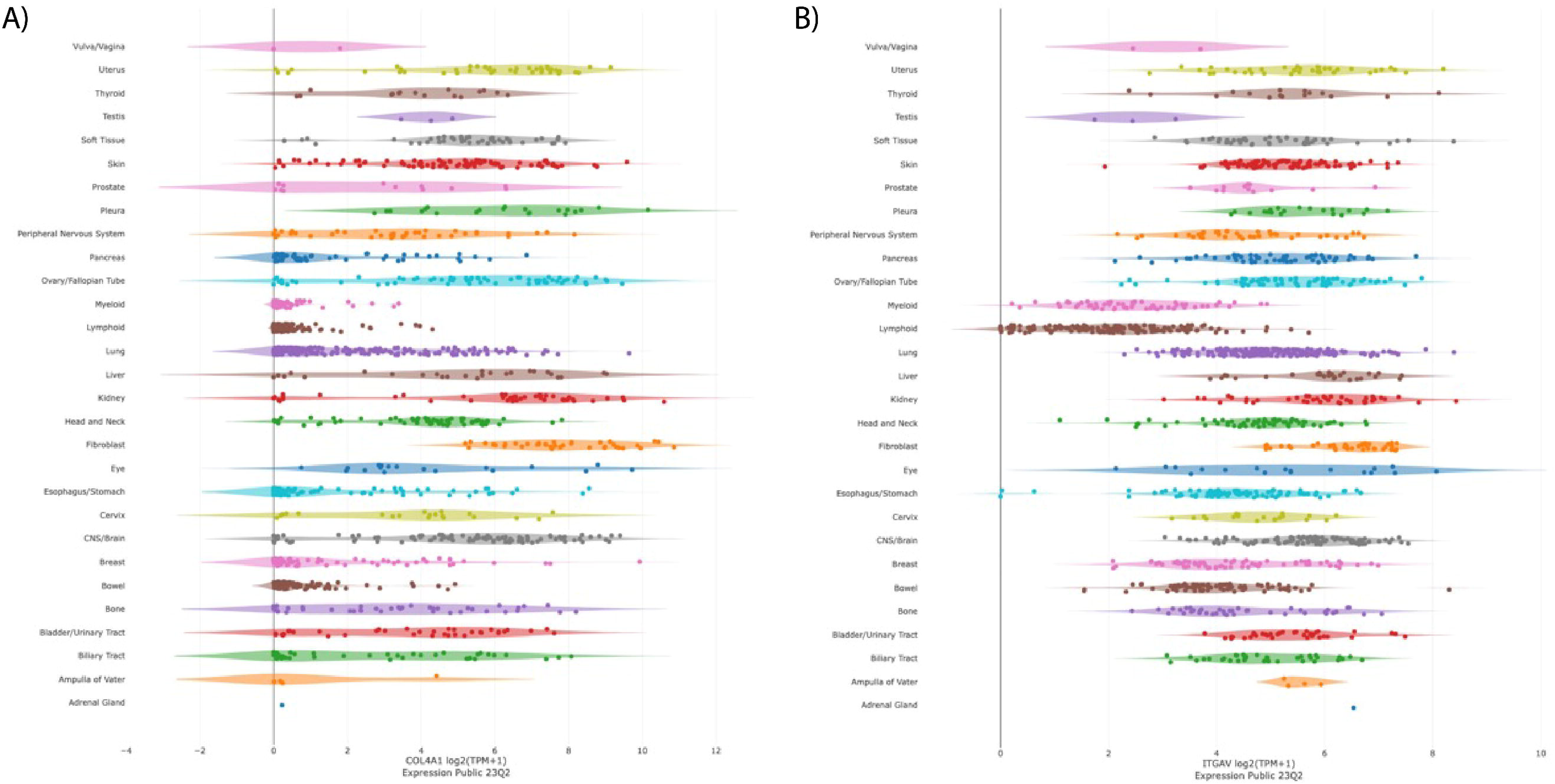
Expression of COL4A1 and ITGAV in normal tissues from Genotype-Tissue Expression (GTEx) project.

Supplementary Table 1: Number of cells that passed QC on primary tumor FOVs.

Supplementary Table 2: Differentially expressed genes identified from subclustering InSituType-assigned T cells and mononucleic phagocytes.

Supplementary Table 3: Differential gene expression using "FindMarkers" between malignant proximal tubule cells and normal proximal tubule. These cells were used for LASSO regression and genes selected by LASSO for final model.

Supplementary Table 4: Abundance differences for cell phenotypes in the tumor and stromal FOVs of the cohorts. Positive estimate indicates lower abundance in IO Naïve non-sarcomatoid.

Supplementary Table 5: Association of Ripley’s K between cohorts in tumor and stromal FOV tissue at a radius of 150px (27um). Negative Estimate values indicate lower clustering in IO naïve non-sarcomatoid samples.

Supplementary Table 6: Differential gene expression of primary ccRCC tumor and stroma samples between cohorts.

Supplementary Table 7: Differential gene expression for each cell phenotype in primary tumor FOVs between pre-and post-IO.

Supplementary Table 8: Differential gene expression in stromal FOVs, between pre-and post-IO primary samples.

Supplementary Table 9: Differential gene expression for each phenotype on primary tumor FOVs before undergoing IO with and without sarcomatoid features.

Supplementary Table 10: Differential gene expression for each phenotype on primary stromal FOVs before undergoing IO with and without sarcomatoid features.

Supplementary Table 11: T-test results comparing bivariate Moran’s I of ligand/receptor pairs from EMT gene set in tumor and stromal FOVs.

Supplementary Table 12: T-test results comparing bivariate Moran’s I of ligand/receptor pairs from IL6/JAK/STAT3 signaling gene set tumor and stromal FOVs.

Supplementary Table 13: Comparisons between ITGAV and COL4A1 abundance in the tumor and stromal compartments.

Supplementary Table 14: Results of beta-binomial modeling of phenotypes between cohorts of the validation mIF.

Supplementary Table 15: Nanostring CosMx SMI probe targets.

Supplementary Table 16: Kidney Cell Atlas phenotype names before (InsituType_Manual) and after (Final) collapsing similar cell types.

