## Supplemental Text for "Increased spatial coupling of integrin and collagen IV in the immunoresistant clear cell renal cell carcinoma tumor microenvironment"

**Pseudo-Bulk Gene Expression in Primary Tissues:**

To assess broadly genes that were differentially expressed in patient cohorts, we collapsed the expression of all cells per FOV to compare impacts of IO treatment. Tumor FOVs showed 61 genes significantly higher in IO naïve samples compared to IO exposed, while IO exposed tumor FOVs showed 9 genes with higher expression compared to IO naïve **(Supplementary Table 6)**. The most significant differentially expressed gene in tumor FOVs was *TPSB2* (FDR = 0.003) which was significantly higher in pre-IO samples. Stromal FOVs also showed more genes having significantly higher expression in IO-naïve (79 genes) than IO exposed tissue samples. The adenosine receptor *ADORA2A*, previously shown to have higher expression in metastatic tissue (31), was not significantly higher in IO naïve compared to IO exposed (tumor FDR = 0.214, stroma FDR = 0.570); however there was a trend for slightly higher expression in both the tumor and stroma FOVs.

**Pseudo-Bulk Differentially Expressed Genes of Sarcomatoid Features**

Taking the average gene expression from all cells within FOVs allows us to assess the difference in gene expression between patient groups, similar to bulk RNA sequencing. We did not observe any genes that were significantly different between those IO naïve primary tumors with and without sarcomatoid features at a threshold of FDR = 0.1 (**Supplementary Table 6**). The gene that showed the most difference in stroma FOVs, *KITLG*, had an FDR = 0.15 and trended towards slightly higher SCT expression in stroma FOVs without sarcomatoid features. The most significantly different gene expression found in tumor FOVs was *MEG3* with an FDR of 0.19, pointing to no differences in global gene expression in the tumor FOVs.

**Ligand-Receptor Autocorrelation in Tissues with Sarcomatoid Features:**

Using bivariate Moran’s I to determine spatial autocorrelation of gene expression belonging to the EMT gene set did not identify differences between tumor FOVs from tissues with and without sarcomatoid features. The largest difference was identified for *THBS1* and *ITGB1* (FDR = 0.129) had a mean Moran’s I of 0.125 in sarcomatoid samples and -0.009 in non-sarcomatoid samples, though not statistically significant (**Supplementary Table 11**). Highest mean Moran’s I in samples with sarcomatoid features was *COL4A1* and *ITGB1* (I = 0.26) followed by several other collagen-*ITGB1* pairs. Gene pairs from the IL6/JAK/STAT gene set ligand-receptor pairs all showed low Moran’s I (highest mean of I = 0.02 for *LTB* and *LTBR* in samples with sarcomatoid features and I = 0.18 for *TNF* and *TNFRSF1B* in IO naïve primary tumors without sarcomatoid features) and high FDR values (FDR > 0.60).

Stromal FOVs also showed no differences between those from tissues with and without sarcomatoid features (**Supplementary Table 11**). Spatial autocorrelation of ligand-receptor gene pair was similar between sample groups with FDR > 0.70. While no difference between groups was observed, mean Moran’s I values for *COL4A1* and *ITGB1* were positive indicating spatial autocorrelation (I = 0.133 in samples with sarcomatoid features and 0.124 in samples without sarcomatoid features), similar to results found in the tumor FOVs. The IL6/JAK/STAT gene set did not show high Moran’s I values, with the largest autocorrelation coming from *TNF* and *TNFRSF1B* in sarcomatoid samples (I = 0.025). This pair was also the most different between tissues with and without sarcomatoid features (FDR = 0.118), and tissues without sarcomatoid features showed negative I = -0.024.
